# Protein response to deleterious episodes of GC-biased gene conversion in mammals of different Ne

**DOI:** 10.64898/2026.10.02.756222

**Authors:** Marie Riffis, Maxime Locatelli, Nathanaëlle Saclier, Nicolas Galtier

## Abstract

GC-biased gene conversion (gBGC) is a widespread recombination-associated process that favours the fixation of GC alleles independently of their fitness. In coding regions, it can oppose purifying selection and favour the accumulation of deleterious AT to GC mutations.In mammals where recombination hotspots are predominantly determined by PRDM9, the deleterious effect of gBGC is expected to be episodic due to the rapid turnover of hotspots. The deleterious impact of gBGC episodes on proteins has been well documented, particularly in rodents and primates. However, the evolutionary response of the affected proteins after the episodes end has been scarcely investigated. Theory predicts that a decline in gBGC intensity should trigger a compensatory response that enables previous damage to be repaired. Recently, we reported evidence for pervasive compensatory evolution following gBGC episodes in *Hydromyini* rodents. Here, we aimed to investigate whether this post-gBGC compensation is widespread among mammals in which gBGC is episodic. Using a phylogenetic approach, we analysed substitution patterns in coding exons from five mammalian taxa spanning a range of effective population size (Ne): *Murini*, *Rattini*, *Cetacea*, *Cercopithecidae* and *Hominoidae*. We detected thousands of gBGC episodes, characterised by an increased rate of non-synonymous AT to GC substitutions.We found evidence for compensatory evolution in non-primate taxa, and a lack of signal in Primates. Although the influence of Ne remains unclear, our study highlights both the evolutionary significance of post-gBGC compensation and the methodological challenges in detecting it.

## Introduction

GC-biased gene conversion (gBGC) is a molecular process that occurs during meiotic recombination and is caused by a bias towards GC bases in DNA repair. Acting like a segregation distorter, gBGC increases the population frequency and the fixation probability of GC alleles, independently of their fitness (Galtier et al., 2001). Over the past few decades, there has been a growing interest in this mechanism, with many studies being conducted, mostly on mammals. These studies have demonstrated that gBGC is widespread in the living world and has a strong influence on the evolution of genomic landscapes in many eukaryotic species (Capra & Pollard, 2011; Duret & Galtier, 2009; Figuet et al., 2015; Galtier et al., 2018; Pessia et al., 2012). Notably, gBGC increases the GC content in highly recombining regions (Duret & Arndt, 2008; Mugal et al., 2015; Pessia et al., 2012) and can confound classical tests of positive selection (Berglund et al., 2009; Bolívar et al., 2016; Corcoran et al., 2017; Dreszer et al., 2007; Galtier & Duret, 2007; Galtier et al., 2009; Ratnakumar et al., 2010; Rousselle et al., 2019).

Importantly, gBGC is expected to counteract purifying selection, thus generating a mutation load within genomes by favouring the fixation of slightly deleterious AT to GC mutations in functional regions located near high-recombining regions. The deleterious effects of gBGC have been demonstrated both theoretically (Duret & Galtier, 2009; Glémin, 2010) and empirically (*e.g.* Berglund et al., 2009; Galtier et al., 2009; Haudry et al., 2008). In mice, the Fxy gene, which was recently translocated into the highly-recombining region of the X chromosome, has undergone the fixation of many deleterious AT to GC substitutions over a very short period of time (Galtier & Duret, 2007). In humans, disease-associated polymorphisms are maintained by gBGC in highly-recombining regions, leading to an increased risk of hereditary diseases (Lachance & Tishkoff, 2014; Necşulea et al., 2011).

This deleterious effect of gBGC on protein-coding genes is expected to be episodic in many mammals, due to the evolutionary instability of the recombination rate at fine genomic scales. At this scale, recombination is concentrated in hotspots, the locations of which are predominantly determined by the PRDM9 protein in most mammals (Baudat et al., 2010; Grey et al., 2018; Myers et al., 2010; Parvanov et al., 2010). PRDM9 and its DNA-binding motifs evolve very rapidly, resulting in a rapid turnover of recombination hotspots (Baker et al., 2015; Berg et al., 2011; Latrille et al., 2017; Lesecque et al., 2014; Myers et al., 2010; Oliver et al., 2009). Several studies have demonstrated the episodic nature of gBGC by analysing the substitution pattern between closely related species. These analyses showed a clustering of AT to GC substitutions in specific lineages and regions, often close to recombination hotspots (Berglund et al., 2009; Capra & Pollard, 2011; Dreszer et al., 2007; Galtier, 2021; Galtier et al., 2009). Berglund et al., 2009 and Galtier et al., 2009 have demonstrated that these gBGC episodes result in an accumulation of non-synonymous substitutions, probably deleterious, in primate proteins.

Although the deleterious impact of gBGC on protein-coding genes is well documented, the evolutionary response of the affected proteins remains largely unexplored. Because the accumulation of deleterious AT to GC mutations is episodic, it has been hypothesised that, once a recombination hotspot dies and the gBGC episode ends, compensatory evolution could bring back the protein to its fitness optimum (Galtier et al., 2009; Riffis et al., 2025). A theoretical study indeed suggested that gBGC episodes could significantly alter the evolutionary trajectory of the affected proteins, with a decrease in gBGC strength being followed by an increase in the fixation rate of beneficial mutations (Joseph, 2024). Recently, we analysed the evolution of coding sequences in *Hydromyini* rodents and found evidence for pervasive compensatory evolution following gBGC episodes in this group (Riffis et al., 2025). Using a phylogenetic approach, we first identified gBGC episodes as clusters of AT to GC synonymous substitutions in specific exons and lineages. Then, we analysed protein evolution in the downstream branches and detected an increased rate of non-synonymous substitutions, likely due to the fixation of beneficial mutations that compensate for the deleterious substitutions caused by the episode.

The report of Riffis et al., 2025 newly describes a process by which proteins recurrently move away from their fitness optimum due to gBGC-induced deleterious mutations, then back towards the optimum via compensatory evolution. This study concerned a single tribe of rodents, and it is currently unclear whether the newly-discovered process is general, or a specificity of this (or a small number of) taxa. This is an important question to answer: is it really the case that proteins recurrently undergo bursts of amino-acid substitutions due to the birth-death of recombination hotspots, regardless of environmental changes? To confirm the existence of this mechanism at a broader taxonomic scale is therefore an important goal. The effect of the effective population size (Ne) would appear particularly interesting to consider. The small-sized rodents analyzed by Riffis et al., 2025 are relatively large-Ne mammals. A large body of literature demonstrates that large, long-lived mammals, such as primates or artiodactyls, typically have smaller Ne’s than rodents, and therefore experience a higher rate of genetic drift and less effective purifying selection (*e.g.* Figuet et al., 2016; James and Eyre-Walker, 2020; Lartillot, 2013a; Popadin et al., 2007; Romiguier et al., 2013.

Ne could in principle influence the process we here consider in two ways. First, Ne might modulate the effect of gBGC. Like selection, gBGC is only effective if stronger than drift. If the gBGC rate, b, was constant, then the efficiency of gBGC, which is determined by the Ne.b product, would be decreased in small-Ne species. Both theoretical and empirical arguments suggest that b is negatively related to Ne, such that Ne.b might be roughly constant among lineages (Clessin et al., 2025; Galtier et al., 2018). However, in mammals the strength of gBGC has been found to be higher in large-Ne rodents or *Chiroptera* than in small-Ne primates or artiodactyls (Galtier, 2021; Lartillot, 2013b; Romiguier et al., 2010). It is therefore unclear whether the deleterious effect of gBGC episodes on proteins reported by Riffis et al., 2025 in *Hydromyini* similarly affects molecular evolution in small-Ne groups of mammals. Secondly, Ne might affect the efficiency of the compensatory response to deleterious episodes.

Adaptation is expected to be more efficient in large populations (Lanfear et al., 2014). Large-Ne species are expected to exhibit greater standing genetic variation and to produce more new mutations per generation than small-Ne ones. Furthermore, slightly beneficial mutations have a higher fixation probability in large than in small populations. Empirical measures of the protein adaptive rate, however, have provided mixed evidence regarding the link between the adaptive rate and Ne: a positive relationship was detected among closely related species (Leroy et al., 2021; Rousselle et al., 2020), but not among more distantly related ones, despite stronger contrasts in Ne (Galtier, 2016). It will be interesting to check whether small-Ne species of mammals re-adapt similarly to large-Ne ones after deleterious gBGC episodes.

The study on *Hydromyini* used a large set of exons and showed that detecting gBGC episodes and subsequent compensation requires high-resolution taxonomic sampling (Riffis et al., 2025). Furthermore, the analysis must be conducted in groups where PRDM9 is at work, ensuring that gBGC is episodic. Here, we analyzed five mammalian groups meeting these criteria : two groups of *Murinae* rodents, *Rattini* and *Murini* (large Ne), and three groups of small-Ne mammals: *Hominoidae* (apes), *Cercopithecidae* (Old World monkeys), and *Cetacea*. Our results indicate that, although the strength of gBGC varies among mammalian taxa, compensatory evolution following deleterious episodes of gBGC is probably a widespread phenomenon.

## Results

### Overview of the dataset diversity

We generated high-quality alignments of coding exons from whole-exome sequencing in *Murini* and *Rattini*, genome annotation in *Hominoidae* and *Cercopithecidae*, and pre-existing exon alignments in *Cetacea*. In each of these taxa, we reconstructed the species phylogeny from the concatenation of exon alignments (Supplementary Figure S1-4). Then for each exon we mapped synonymous (S) and non-synonymous (NS) nucleotide substitutions to the phylogeny, distinguishing 3 types of substitutions: AT to GC (Weak to Strong: WS), GC to AT (Strong to Weak: SW) and GC-conservative (SSWW). The number of substitutions mapped for each type is detailed for each taxon in Supplementary Table 1. The synonymous substitution pattern for each taxon (Figure 1) shows that rodents have more branches with more than one S substitution. Furthermore, S-WS substitutions tend to cluster within branches, as shown by their increasing proportion with the total number of substitutions in a branch, for rodents and *Cetacea* but not for Primates (Figure 1). The taxa used for the analyses differ in terms of divergence time, phylogenetic sampling, number of exons and substitutions, and tree branch lengths (Table 1). In particular, mean branch length is two to four times shorter in *Cetacea* and Primates than in the rodent data sets. Furthermore, *Cetacea* and Primates have a lower total number of substitutions relative to the overall amount of data (i.e. number of species and exons) than the other groups. The distributions of exon length, branch length, GC3 and GC12 are shown in Supplementary Figure S6A-10A. All distributions are quite similar between groups, except the GC3 distribution that is much tighter in rodents.

**Figure 1:**
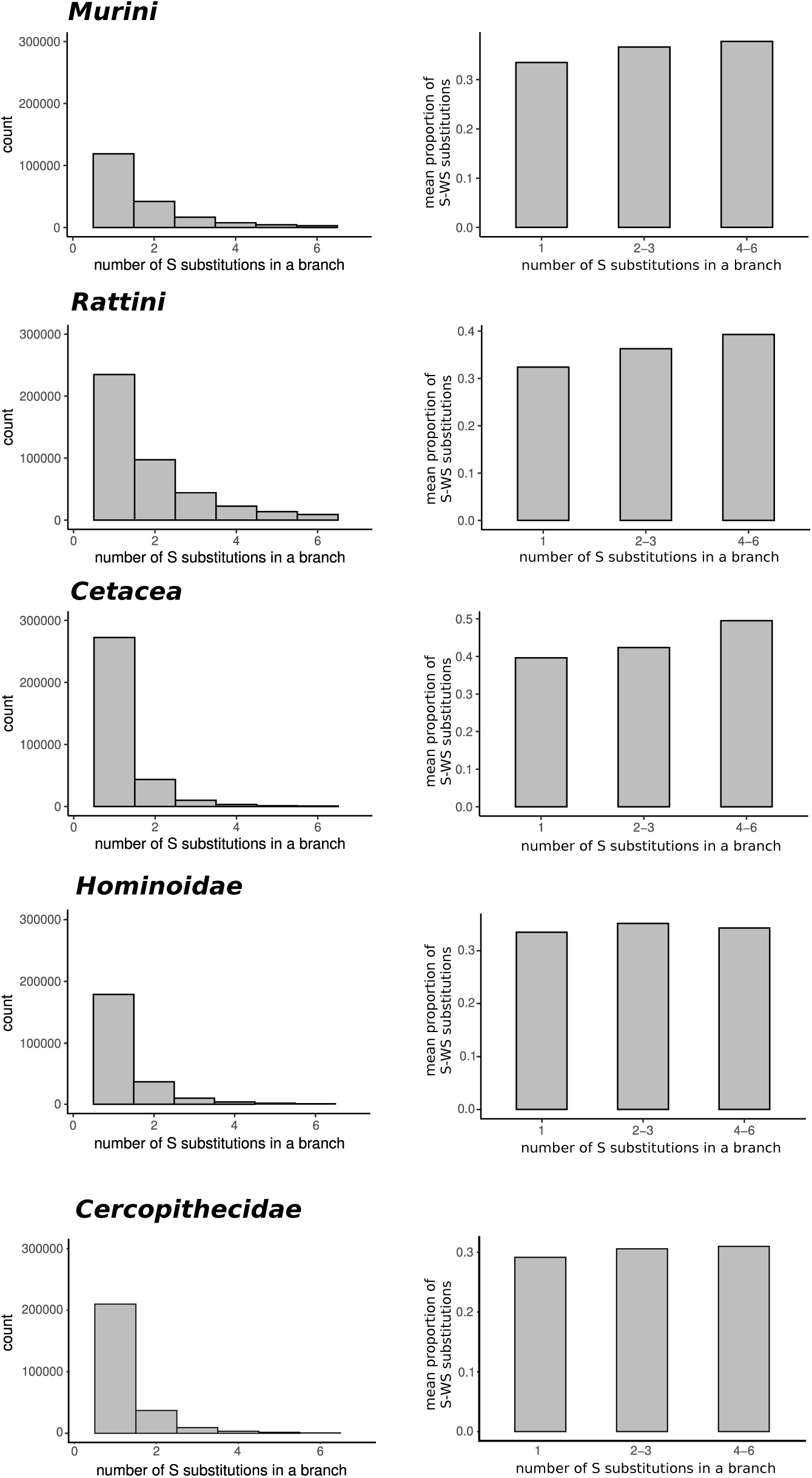
Pattern of synonymous substitutions in the different taxa. Left side: distribution of the number of S (all types) substitutions in branches. Right side: clustering of the S-WS substitutions in the tree. For both sides, data for branches with 0 and with more than 6 S substitutions are not shown.

**Table 1:**
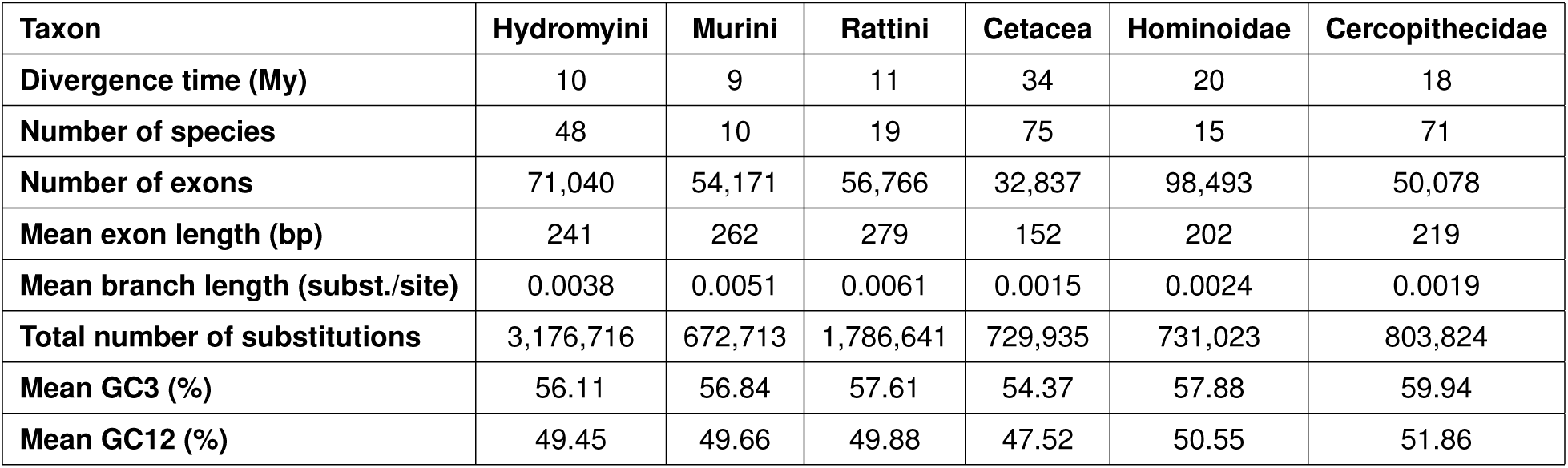
Summary of the main features of the different taxa studied. For comparison, we included the data on Hydromyini from Riffis et al. (2025). Reference for the divergence times: https://timetree.org/

### Detection of the gBGC episodes

We first identified gBGC episodes, which are characterised by a clustering of synonymous WS (S-WS) substitutions across branches and exons. A gBGC episode is detected in a specific branch and exon if there is an excess of S-WS substitutions compared to the expectation under a random distribution of substitutions across branches. Using this method, thousands of episodes were detected in each taxon: 3,301 in *Murini*, 14,580 in *Rattini*, 9,999 in *Cetacea*, 2,978 in *Hominoidae* and 4,245 in *Cercopithecidae*. Groups with a higher total number of substitutions had a higher number of detected episodes (Table 1). The distributions of exon length, branch length, GC3 and GC12 for the detected episodes are shown in Supplementary Figures S6B-S10B. Overall, the distributions are almost identical to those in the full dataset, but the exons and branches are slightly longer on average.

Episodes occurring in internal branches were then classified into three categories: punctual (episodes occurring in a single branch and exon), two-branch (episodes occurring in a single exon, but in the focus branch and the downstream branches), and all-exons (gene-wide episodes) (Table 2). The classifier is based on an ML method that provides the posterior probability of each episode belonging to a category (see the supplementary material of Riffis et al., 2025 for details). We assigned an episode to a category if its posterior probability of belonging to that category was higher than the posterior probabilities of belonging to the other two categories. For *Cetacea*, only two categories could be defined (see Materials & Methods): punctual and non-punctual, and an episode was classified as punctual if its posterior probability of belonging to this category was superior to 0.7. For all taxa, the majority of episodes were classified as punctual (Table 2). Only the punctual episodes were kept for subsequent analysis, since they are localised and short-lived, as expected from episodic gBGC resulting from short-lived recombination hotspots driven by PRDM9.

**Table 2:** Details of the number of gBGC episodes detected in each taxon. The “total number of episodes” includes episodes in all branches of the tree, whereas the three episode types (“punctual”, “two-branch”, “all-exons”) include only episodes found in internal branches.

| Taxon | All episodes | Punctual episodes |  | Two-branch episodes | All-exons episodes |
| --- | --- | --- | --- | --- | --- |
|  |  | Deleterious | Non-deleterious |  |  |
| **Murini** | 3,301 | 170 | 686 | 0 | 8 |
| **Rattini** | 14,580 | 696 | 1,927 | 329 | 1 |
| **Cetacea** | 9,999 | 732 | 2268 | - | - |
| **Hominoidae** | 2,978 | 658 |  | 47 | 7 |
| **Cercopithecidae** | 4,245 | 894 |  | 51 | 45 |

As expected, we observed many more S-WS substitutions than other types in punctual episodes, compared to branches without episodes (Supplementary Table 1). This pattern was not detected in branches after punctual episodes (Supplementary Table 1), suggesting that the sorting of the episodes has efficiently selected localised, short-lived episodes that do not overflow into the downstream branches.

An excess of non-synonymous WS (NS-WS) substitutions relative to NS-SW was observed in branches with punctual episodes, but not in branches without episodes, in rodents and *Cetacea* (Figure 2). This suggests that gBGC episodes increase the rate of NS-WS substitutions, in agreement with the suggestion that gBGC episodes have a deleterious effect by promoting the fixation of undesired WS changes. Such a pattern was not observed in the two primates data sets, suggesting that the deleterious effect of gBGC episodes is not dominant in these groups. Given this, and the fact that we did not observe an overall significant clustering of S-WS substitutions in this group (Figure 1), we did not investigate post-gBGC compensation in these taxa. Punctual episodes were then separated into two categories for subsequent analysis (Table 2). The first category included episodes associated with at least one NS-WS substitution, which is supposedly deleterious and has potentially impacted the protein fitness. We called these “deleterious episodes”. The second category includes episodes that were not associated with any NS-WS substitution. We called these “non-deleterious” episodes. Compensatory evolution, if any, is expected to only occur after deleterious episodes.

**Figure 2:**
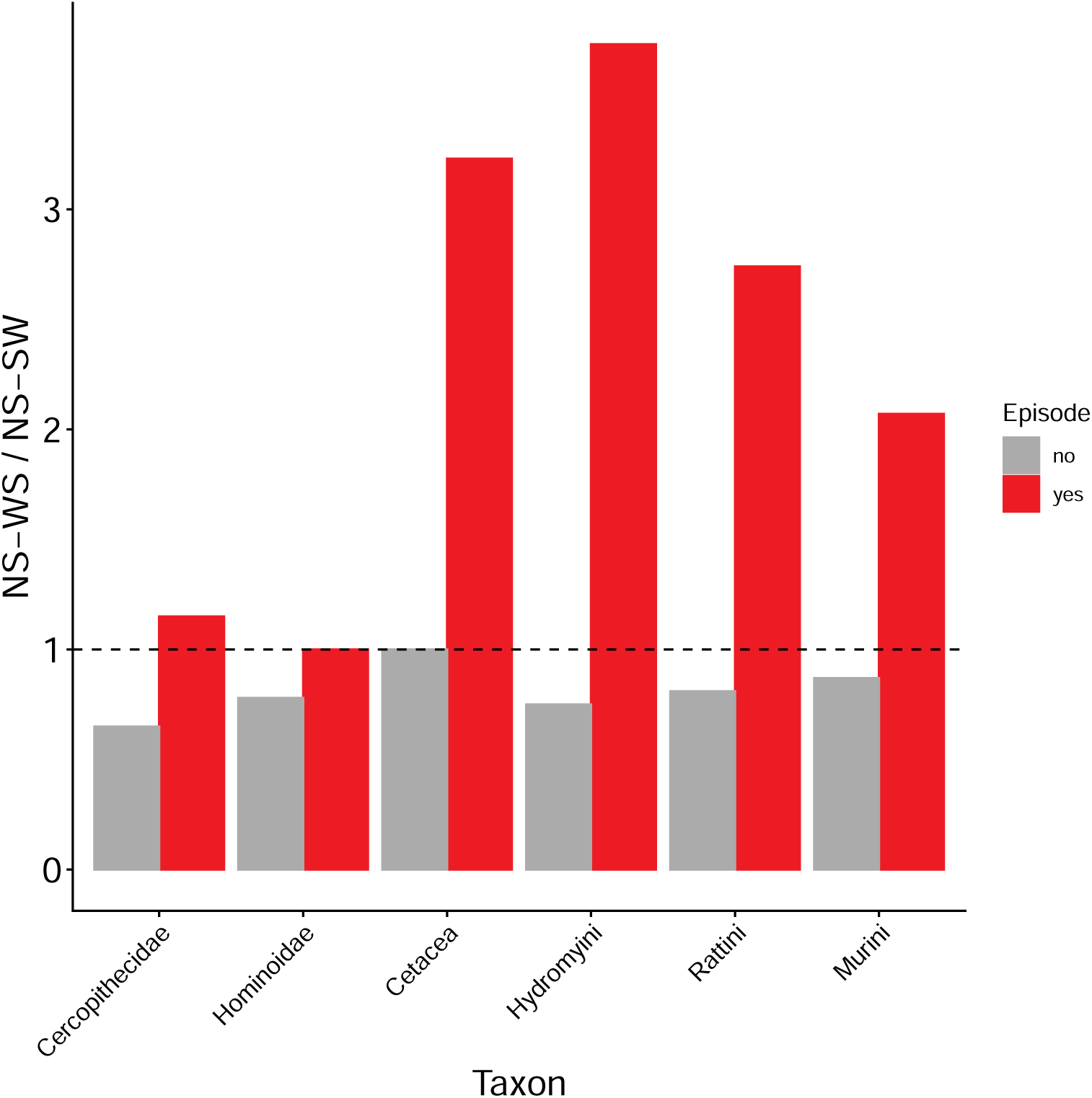
Non-synonymous substitution pattern in the detected gBGC episodes. The y axis represent the ratio of NS-WS over NS-SW substitutions in branches with (right column, in red) and without episodes (left column, in grey). The Hydromyini data come from Riffis et al. (2025).

### Protein evolution after punctual gBGC episodes

We then looked for a compensatory response following punctual gBGC episodes, distinguishing deleterious ones from non-deleterious ones. We compared the number of NS substitutions in the branches downstream of the episodes with an expected distribution generated by randomising NS substitutions across exon trees. The rationale behind this is that a significant excess of NS substitutions after deleterious punctual episodes but not after non-deleterious ones is interpreted as reflecting the fixation of compensatory mutations in response to the deleterious effects of gBGC episodes. Our initial analyses of compensatory evolution revealed a weak signal of compensation following punctual episodes across taxa (Figure 3). We observed a non-significant excess of NS substitutions after deleterious episodes in *Murini* and *Rattini*. In *Cetacea*, we observed almost no excess after either deleterious or non-deleterious episodes.

**Figure 3:**
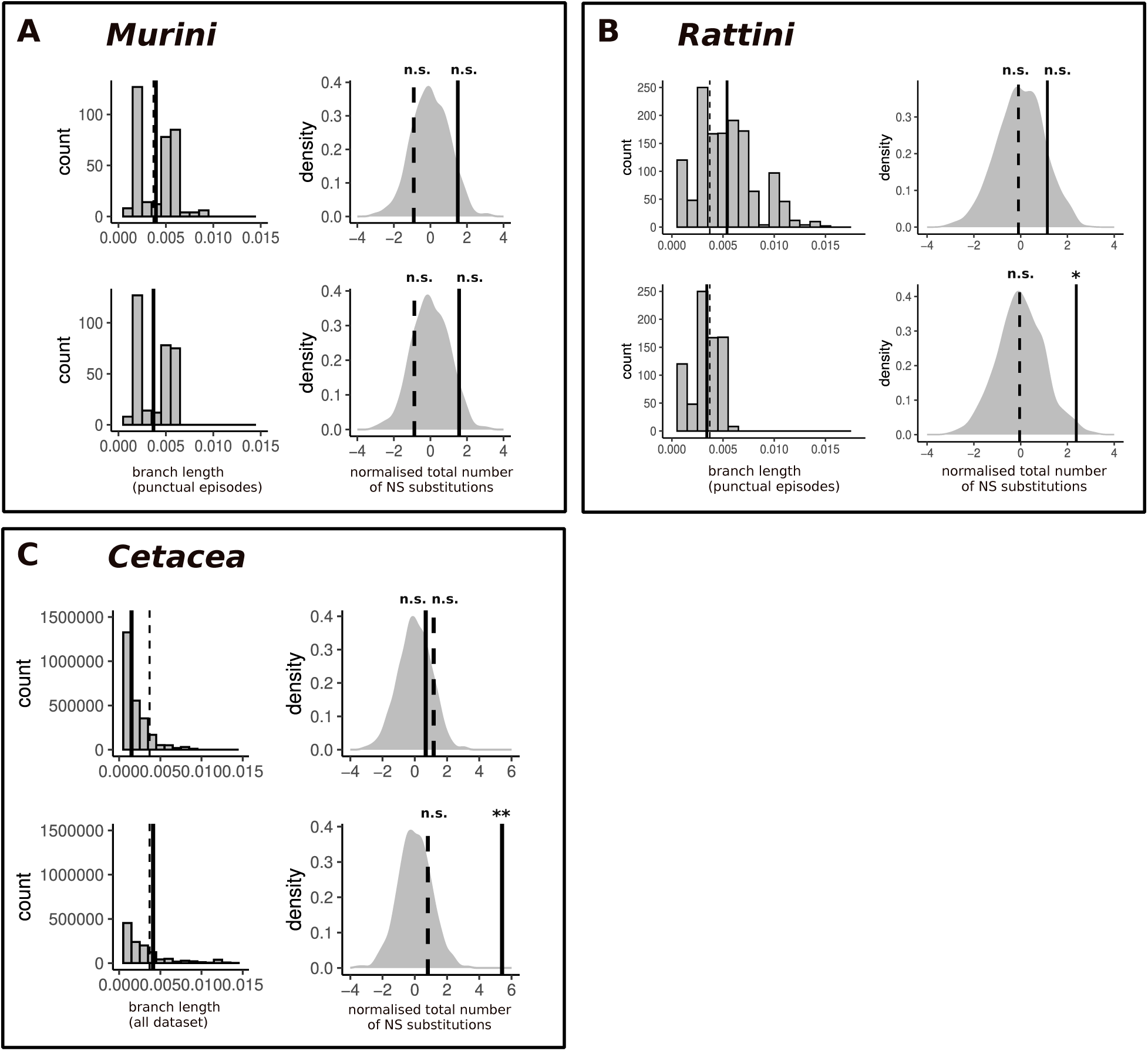
Protein evolution after punctual gBGC episodes and effect of adjusting the branch lengths. A: Murini and B: Rattini, for which we adjust the length of the branches carrying the episodes, by applying a filter of length *<* 0.006 substitution per site. C: Cetacea, for which we adjust the length of the branches of all the dataset, by subsampling the tree to have a maximum of 25 species. The left side shows the distribution of the branch lengths, with the vertical solid line corresponding to the mean for the given taxon, and the vertical dashed line corresponds to the mean for the Hydromyini dataset. The right side shows the total number of NS substitutions after punctual episodes, which was standardised by subtracting the mean and dividing by the standard deviation of the distribution. The distribution shows the expected numbers from 1000 randomisations. The vertical solid line corresponds to the observed number after deleterious episodes and the vertical dashed line corresponds to the observed number after non-deleterious episodes. n.s. : p-value *≥* 0.05, * : p-value *<* 0.05, ** : p-value *<* 0.01.

Our previous analysis in *Hydromyini* suggested that the length of episode-carrying branches is an important determinant of compensatory evolution detection (Riffis et al., 2025). Indeed, we found that compensation in *Hydromyini* was only observed when the branches carrying the episodes were short enough, and that the signal fades as the branch length increases. As the groups studied here differ a lot in terms of branch length distribution, we conducted additional analyses controlling for this parameter. In *Rattini* and *Murini*, which have a poor phylogenetic sampling and therefore a higher mean branch length than *Hydromyini* (Table 1), we focused on episodes carried by a branch of length inferior to 0.006 substitutions per site, as we did in *Hydromyini*. This reduced the number of deleterious punctual episodes to 158 in *Murini* and to 395 in *Rattini*, and the number of non-deleterious punctual episodes to 633 in *Murini* and to 1,260 in *Rattini*. This enabled us to detect a strong compensation signal in *Rattini*, with a stronger, significant excess of NS substitutions after deleterious episodes that occurred in branches shorter than 0.006 substitutions per site (Figure 3). As in *Hydromyini*, this signal is lost when the episodes branch lengths are longer than this threshold (Supplementary Figure S11). On average, the branch lengths in *Cetacea* are more than twice as short as those of the other groups, due to an exhaustive phylogenetic sampling. We therefore performed subsampling of the data by randomly removing species from alignments to ensure a maximum of 25 species. This enabled us to increase the tree branch lengths without significantly altering the topology or skewing the substitution pattern. After subsampling, the mean branch length increased to 0.0041 substitutions per site, approaching that of our rodent data sets. The number of punctual episodes decreased to 465 deleterious and 1,266 non-deleterious. With the subsampled dataset, we observed a strong, highly significant excess of NS substitutions after deleterious episodes but not after the non-deleterious ones (Figure 3C).

As in *Hydromyini*, the compensatory response observed in *Rattini* and *Cetacea* is primarily driven by WS substitutions, and also by SSWW substitutions in *Cetacea* (Supplementary Figure 12). The same is true for *Murini*, for which there is a significant excess of NS-WS substitutions after deleterious episodes, even if the response is not significant when all NS substitutions are considered (Supplementary Figure 12).

## Discussion

We investigated whether the evidence for compensatory evolution following gBGC episodes found in *Hydromyini* rodents, as reported in Riffis et al., 2025, could be extended to other mammalian groups and particularly how the response might vary according to the effective population size. Applying the same method of analysing the substitution pattern in exon coding sequences, we identified exon-specific and lineage-specific gBGC episodes in *Murini*, *Rattini*, *Cetacea*, *Hominoidae* and *Cercopithecidae.* For three of these groups, the branches carrying the episodes exhibited increased rates of NS substitutions, which are likely to be deleterious. We found evidence of compensatory evolution in branches downstream of episodes in rodents and *Cetacea*.

### Deleterious gBGC episodes in rodents and *Cetacea* but not in primates

In *Murinae* rodents, the ratio of gBGC episodes over the number of analysed branches was overall more important than in other groups (Tables 1 & 2), and the episodes were associated with an increased rate of NS-WS substitutions (Supplementary Table 1). This is not surprising, given that previous studies demonstrated strong gBGC in rodents (Clement & Arndt, 2011; Galtier, 2021; Pracana et al., 2020). In *Hydromyini*, a significant proportion of the episodes were classified as “all-exons”, indicating that entire genes, and perhaps even larger regions, undergo prolonged periods of GC-enrichment. We found almost no such type of gBGC events in *Murini* and *Rattini*, consistent with the results of Pracana et al., 2020, who reported the existence of such large scale events of GC-enrichement in the *Mus musculus* (*Murini*) and *Rattus norvegicus* (*Rattini*) lineages.

The detection of thousands of gBGC episodes in *Cetacea*, despite their low substitution rate, is an unexpected and original result since gBGC has not been extensively studied in this group so far. One study found a gBGC strength much lower than in rodents and much higher than in primates in dolphin *Tursiops truncatus* (Lartillot, 2013b), consistent with the report by Romiguier et al., 2010 of an intermediate level of GC3 in *T. truncatus* compared to primates and rodents. Together, these two studies suggest important levels of gBGC in this group, despite only one *Delphinidae* species being examined across all mammals. Our analyses, based on nearly exhaustive sampling of *Cetacea*, confirm these results. Like in murine rodents, the detected gBGC episodes were in *Cetacea* associated with a burst of NS-WS substitution, suggesting a (probably negative) impact on protein evolution (Figure 2).

In primates, we detected no clustering of S-WS substitutions (Figure 1), a lower number of significant episodes (Table 2) than in other groups, and no evidence of an effect of episodes on protein evolution (Figure 2). These observations are in line with the existing literature on gBGC in hominid primate coding sequences. Galtier, 2021 found almost no clustering of the GC substitutions in *Hominidae* and a weak clustering in *Cercopithecidae*, in contrast to a high clustering in *Murinae*. Lartillot, 2013b demonstrated that *Hominoidae* are among the mammalian groups with the weakest gBGC strength (4Neb *<* 1). Galtier et al., 2009 identified only about thirty episodes of accelerated protein evolution with a significant bias toward GC (on 4 analysed branches) in primates. They demonstrated that the dN/dS ratio increased in accelerated branches, suggesting an accumulation of deleterious mutations; however, this observation was limited to only 9 exons with a high amino-acid substitution rate (see also Berglund et al., 2009). Here we rather defined gBGC episodes based on synonymous changes, irrespective of the non-synonymous substitution rate, and detected no significant effect on protein evolution overall. We suggest that the deleterious effect of episodic gBGC in primates is weak and hard to detect by global approaches such as the one we adopted here.

As in *Hydromyini*, we observed here an increase in the rate of SW and SSWW substitutions in gBGC episodes, albeit much weaker than for WS (Supplementary table 1). Surprisingly, this was particularly marked in *Cetacae*. Riffis et al., 2025 suggested that this increase in substitution rates could be explained by a mutagenic effect of recombination (Halldorsson et al., 2019) or by the variance of the coalescent between loci, creating elevated substitution counts of all types in particularly long branches (Rannala et al., 2020). The reasons why these effects would be more pronounced in *Cetacea* are unclear.

### Compensation signal after deleterious gBGC episodes

In *Rattini* and *Cetacea*, we observed the same pattern of post-gBGC compensation as in *Hydromyini* : an increase in the rate of NS substitutions after punctual episodes having at least one NS-WS substitution (potentially deleterious), but not after those that have not (non-deleterious) (Figure 3B & C). We interpreted this pattern as the fixation of beneficial mutations at other sites that compensate for the deleterious effects of gBGC episodes. The classification of the episodes to select only punctual ones ensured that the observed excess of NS substitutions was indeed a response to gBGC episodes and not just an overflow of episodes. This is confirmed by the rates of S-WS and S-SW substitutions that are similar in the descending branches of punctual episodes (Supplementary Table 1), and by the absence of an excess of NS substitutions after non-deleterious episodes (Figure 3B & C), contrary to what would be expected if there were an overflow of episodes in the descending branches. In *Cetacea*, this argument is reinforced by the observation of a response on GC-conservative substitutions too (Supplementary Figure S12).

Overall, the signal was not significant in *Murini* (Figure 3A), but it was similar to those observed in *Hydromyini*, *Rattini* and *Cetacea*. There was a significant excess of NS-WS substitutions after deleterious episodes in this group (Supplementary Figure S12). The weaker signal observed in *Murini* may be due to a lack of statistical power resulting from the small size of the dataset compared to those of other taxa: there are only 10 *Murini* species and the total number of substitutions is substantially lower than in the two other groups of rodents. It would be interesting to see if the response is stronger with a larger sample size.

### No clear effect of Ne

The results obtained here make it difficult to draw clear conclusions about the effect of Ne on the protein response to gBGC. For large Ne taxa (*Hydromyini*, *Murini* and *Rattini*), we found evidence of homogeneous post-gBGC compensatory evolution across groups. For small Ne taxa, however, we found a strong signal, similar to that observed in rodents, in *Cetacea*, and a lack of signal in primates, in which deleterious gBGC is probably undetectable by our approach. The contrast between *Cetacea* and Primates is surprising, given their similar biological features (Romiguier et al., 2014). On the other hand, our data revealed differences in substitution patterns and gBGC strength between the two groups. Consistent with previous studies (Lartillot, 2013b; Romiguier et al., 2010), our results suggest that gBGC may be stronger in *Cetacea* than in Primates, the former being characterised by a higher clustering of S-WS substitutions, a greater number of detected episodes, and a substantially higher rate of NS-WS substitutions within these episodes.

To observe a pattern of re-adaptation as strong in *Cetacea* as in rodents is somehow surprising. Indeed, it is widely accepted that small Ne taxa adapt less efficiently than large Ne taxa. For a protein to adapt (or re-adapt), beneficial mutations must occur (either through new mutations or existing genetic variation) and become fixed. However, the mutation rate, genetic variation, and the probability of fixation of beneficial mutations depend on Ne. Larger populations have a higher mutation rate (Bell, 2013; Cutter et al., 2013; Lanfear et al., 2014; Sung et al., 2012), so large Ne taxa are more likely to experience the appearance of beneficial mutations. Furthermore, it has been demonstrated that the rate of beneficial mutations limits adaptation in small Ne taxa (Rousselle et al., 2019). Genetic diversity is greater in large populations (Bell, 2013; Kimura et al., 1963; Petit & Barbadilla, 2009; Romiguier et al., 2014; Sung et al., 2012), so large Ne taxa are more likely to already possess beneficial compensatory alleles. Finally, weakly beneficial mutations are more likely to reach fixation in large populations, due to a weaker genetic drift (Lanfear et al., 2014). While the literature reviewed above suggests that large populations may adapt more effectively than small ones, there is currently no clear empirical evidence that the adaptive substitution rate is higher in large Ne taxa. On one hand, some studies have found a higher adaptive substitution rate in large populations than in small ones, in various Eukaryotes species (Gossmann et al., 2012; Jensen & Bachtrog, 2011; Phifer-Rixey et al., 2012; Strasburg et al., 2011). On the other hand, a study of 2 species of Drosophila with a 5-fold difference in Ne found that Ne is not a major determinant of the rate of protein adaptation (Bachtrog, 2008). Furthermore, an empirical study found no impact of Ne on the adaptive substitution rate across 44 Metazoans species (Galtier, 2016), a result corroborated by theoretical studies (Lourenço et al., 2013; Razeto-Barry et al., 2012). Our results, *i.e.* a similarly increased NS substitution rate after gBGC episodes in rodents and *Cetacea,* are consistent with the lack of a strong, consistent relationship between Ne and the adaptive rate in the literature. One possible explanation for our results is that post-gBGC compensatory evolution could affect the amino-acid substitution rate independently of Ne, but that small Ne species may re-adapt less efficiently by overall fixing less favorable mutations than large Ne species. Simulations under Fisher’s geometrical model showed that large populations adapt more efficiently than small populations by making larger adaptive steps but not more steps (Lourenço et al., 2013; Razeto-Barry et al., 2012). These concerns about the fitness effect of the presumably compensatory substitutions could not be captured by our analyses, which only counts substitutions.

However, the contrasting results concerning the small Ne taxa are insufficient to draw general conclusions about the effect of Ne on post-gBGC compensatory evolution. We should analyse other small Ne groups, with stronger gBGC strength than in *Hominoidae*, to see if *Cetacea* constitutes an exception.

### Detectability of the compensation signal

Our analyses demonstrated that the detection of a post-gBGC compensation signal depends heavily on the mean branch length of the dataset (Figure 3). In *Rattini*, we detected a better signal with shorter episode branch lengths and, as in *Hydromyini*, the signal becomes undetectable when the branches carrying the episodes are too long (Supplementary Figure S11). This may be because, if the branches are too long, gBGC episodes and the compensatory response occur within the same branches, meaning that we cannot detect anything in the descending branches (Figure 4). Conversely, the dense phylogenetic sampling in *Cetacea* produced very short branches, and we only detected the compensation signal after subsampling the tree, thus obtaining longer branch lengths. We hypothesise that, in case of a high density of lineages, the compensation signal might be distributed across multiple branches after the episodes, making compensation undetectable when only the two branches immediately downstream of the episode are considered (Figure 4). Additionally, we demonstrated that, even with ‘appropriate’ branch lengths, the compensation signal can be weak, as observed in *Murini* (Figure 3A). We suggest that this is due to a lack of data, as the number of species and total number of substitutions are low in this group, which probably limits the statistical power of our analysis. These observations highlight a methodological limitation of our approach, which requires a large enough number of substitutions and appropriate phylogenetic resolution to reach its maximal power.

**Figure 4:**
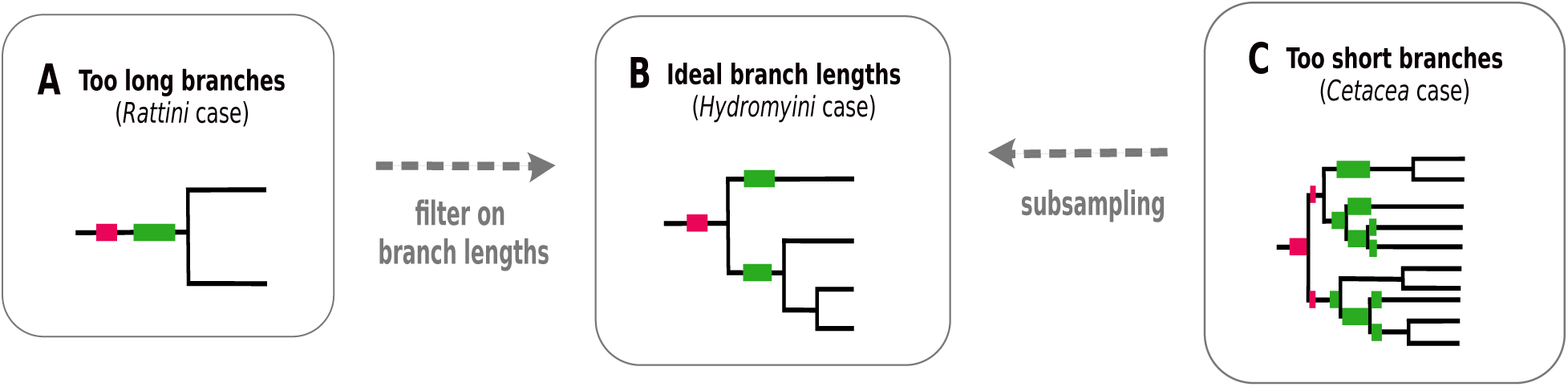
Supposing effect of the phylogenetic resolution on the detection of a compensation signal. A: if the branches are too long, episodes and compensation may occur in the same branch, making it impossible to detect compensation in the descending branches. B: if the branches have the ‘appropriate’ length, episodes can be detected in a branch and compensation in the descending branches. C: if the branches are too short, episodes could overflow into the descending branches and become mixed with compensation. Compensation may extend into several descending branches, such that the signal is diluted and undetectable in the two direct descending branches.

## Material & Methods

### Data

The data came from various sources, depending on the taxon. For *Murini* and *Rattini*, we downloaded whole-exome sequencing raw reads from the NCBI database, generated by Roycroft et al., 2021 for *Rattini* and by Sarver et al., 2017 for *Murini* (references: PRJNA323493 and PRJNA705792, Supplementary Table S2). There were 10 *Murini* species and 19 *Rattini* species. For *Murini*, reads came from two individuals for nine species and from six individuals for one species, and for *Rattini*, they came from one individual for all species. For *Cetacea*, we obtained exon alignments for 75 species from McGowen et al., 2020. For Primates, we downloaded the complete genomes of 15 species of *Hominoidae* and 71 species of *Cercopithecidae* from Ensembl, NCBI and ENA databases, which were generated by Shao et al., 2023 for the NCBI genomes (reference: PRJNA785018) and by Kuderna et al., 2024 for the ENA genomes (reference: PRJEB67744, Supplementary Table S1), as well as genome annotation of *Homo sapiens* and *Macaca mulatta* from the Ensembl database.

### Exon alignments

We applied the same pipeline to *Murini* and *Rattini* as described by Riffis et al., 2025. Briefly, we cleaned and pooled the reads before assembling them using TRINITY v.2.13.2 (Grabherr et al., 2011). We identified orthologous exons using reciprocal BLAST between the obtained contigs and the reference coding exons from the house mouse (downloaded from the Ensembl database). We aligned exon sequences using MACSE (Ranwez et al., 2018), a codon-aware nucleotide alignment tool. We applied filters to the exon alignments, based on heterozygosity and average coverage depth, to exclude potential sequencing errors or paralogy. We also excluded sequences containing stop codons or fewer than 100 bp from the alignments. Finally, we obtained 54,171 and 56,767 exon alignments for *Murini* and *Rattini* respectively. For *Cetacea*, we used existing exon alignments, but performed additional cleaning to retrieve stop codons and excluded alignments of fewer than 50 bp. We also re-aligned the sequences using MACSE to minimise alignment errors and ensure the reading framework was respected.

For Primates, we used the LiftOn software (Chao et al., 2025) to transfer the annotation of the orthologous coding exons from a reference genome (*Homo sapiens* for *Hominoidae* and *Macaca mulatta* for *Cercopithacidae*) to each genome. We then retrieved exon sequences and aligned them with MACSE. We excluded sequences of less than 100 bp and sequences containing stop codons.

Once the exon alignments had been obtained, we applied the same pipeline as Riffis et al., 2025, the steps of which are summarised below.

### Phylogeny and substitution mapping

For each taxon, we reconstructed a species tree using IQ-TREE v.2.0.7 (Minh et al., 2020) from the exons concatenation, using a GTR model and 1,000 ultrafast bootstrap replicates. For rodents and primates, we retained, for each gene, the species for which a sequence of each exon of the gene exists. We then created a tree for each gene by pruning the species tree with the Ape “prune_tree” function. Exons of the same gene therefore have the same topology. For *Cetacea*, we did not require the same set of species for each exon of the same gene due to a lack of data. We therefore created one tree per exon, also by pruning the species tree.

This assumes that the species tree accurately represents the topology of the exons, which may not be the case if there is tree discordance, in particular if exon alignments contain paralogous sequences. We therefore applied a filter to discard exon alignments with potential tree discordance. For each exon, we reconstructed a de novo tree using IQ-TREE. We also used IQ-TREE to constrain the species tree topology and re-estimate the branch lengths according to the alignment. For these two trees, we then calculated the sum of the branch lengths, and discarded the exon alignments for which the ratio of this sum between the constrained tree and the de novo tree exceeded a given threshold. This threshold was set to 1.5 for *Murini*, *Cetacea* and *Rattini*, and to 1.1 for primates. The rationale behind this is that if an alignment does not fit to the species tree, numerous homoplasic substitutions should be mapped when the tree is constrained, resulting in a higher sum of branch lengths than with the de novo tree.

For each taxon, we performed a substitution mapping for each exon alignment. The branch lengths of the tree were optimised under the YN98 codon model (Yang & Nielsen, 1998) with BppML v.3.0.0 (Guéguen et al., 2013). Then, the number of S and NS substitution was estimated in each branch at each codon for the four mutation types (WS,SW,SS,WW) in an empirical Bayesian framework, using MapNH v.2 (Romiguier et al., 2012).

### Detection of gBGC episodes and selection of ‘punctual’ episodes

For each taxon, we identified gBGC episodes in specific branches and specific exons by identifying clusters of S-WS substitutions over time and space. For each exon and branch, the total number of S-WS substitutions was counted and compared to the expected distribution derived from 1000 permutations. These permutations consisted of redistributing the substitutions across the tree, accounting for branch lengths. A gBGC episode was detected if the count was in the top 1% of the expected distribution.

We then used a maximum likelihood classifier to distinguish localised, short-lived gBGC episodes from large-scale and/or long-lasting gBGC events. Using the S-WS and S-SW substitution patterns, the gBGC episodes occurring in the internal branches of the tree were categorised into 3 categories: ‘punctual’, which occurred in a single exon and branch, ‘two-branches’, which occurred in a single exon but extended to the descending branches, and ‘all-exons’, which corresponded to gene-wide gBGC. The model calculated episode-specific posterior probabilities to belong to each category, and a Goodness of Fit (GoF) statistic. We discarded episodes with a GoF inferior to 0.2, and attributed an episode to a category if its posterior probability of belonging to this category was higher than its probability of belonging to other categories. Only ‘punctual’ episodes were of interest for our subsequent compensation analyses. For *Cetacea*, we were only able to distinguish two categories: ‘punctual’ and ’non-punctual’ episodes, because the different topologies of exons within the same gene make it impossible to compare substitution patterns between the same branches across all exons belonging to the same gene. Furthermore, the criteria for defining an episode as ‘punctual’ differed slightly for this group, we required a posterior probability greater than 0.7.

### Detection of post-gBGC compensatory evolution

To test for compensatory evolution, we examined NS substitutions in the lineages downstream of the punctual episodes. We compared the total number of observed NS substitutions with the expected distribution obtained from 1,000 permutations in which substitutions were redistributed across the tree, taking into account the branch lengths and blocking the branches carrying episodes. The observed values and the expected distributions were summed across all branches immediately downstream of punctual episodes in all affected exons. We distinguished two types of punctual episode: those associated with at least 1 potentially deleterious NS-WS substitution (which we called ‘deleterious episodes’, and those not associated with any NS-WS substitution (which we called ‘non-deleterious episodes’). Non-deleterious episodes therefore served as a control, such as if an excess of NS substitutions was observed after deleterious episodes but not after non-deleterious ones, this was interpreted as a compensatory response to the potentially deleterious effect of gBGC episodes, occurring via the fixation of NS compensatory mutations.

## Data accessibility

The scripts used to apply the same pipeline as Riffis et al., 2025 were already available from https://github.com/Marie-98/gBGC_comp, to which we added the scripts to make the annotation transfer for primates. Exon alignments and analysed data files are available from https://doi.org/10.5281/zenodo.17401939.

## Acknowledgements

We would like to thank Michael R. McGowen for sharing the Cetacea exon alignments with us, and Johanna Girodolle, Nathalie Chantret, Vincent Ranwez and Céline Scornavacca for their advice and assistance with the annotation transfer.

