## supplementaries for "Protein response to deleterious episodes of GC-biased gene conversion in mammals of different Ne"

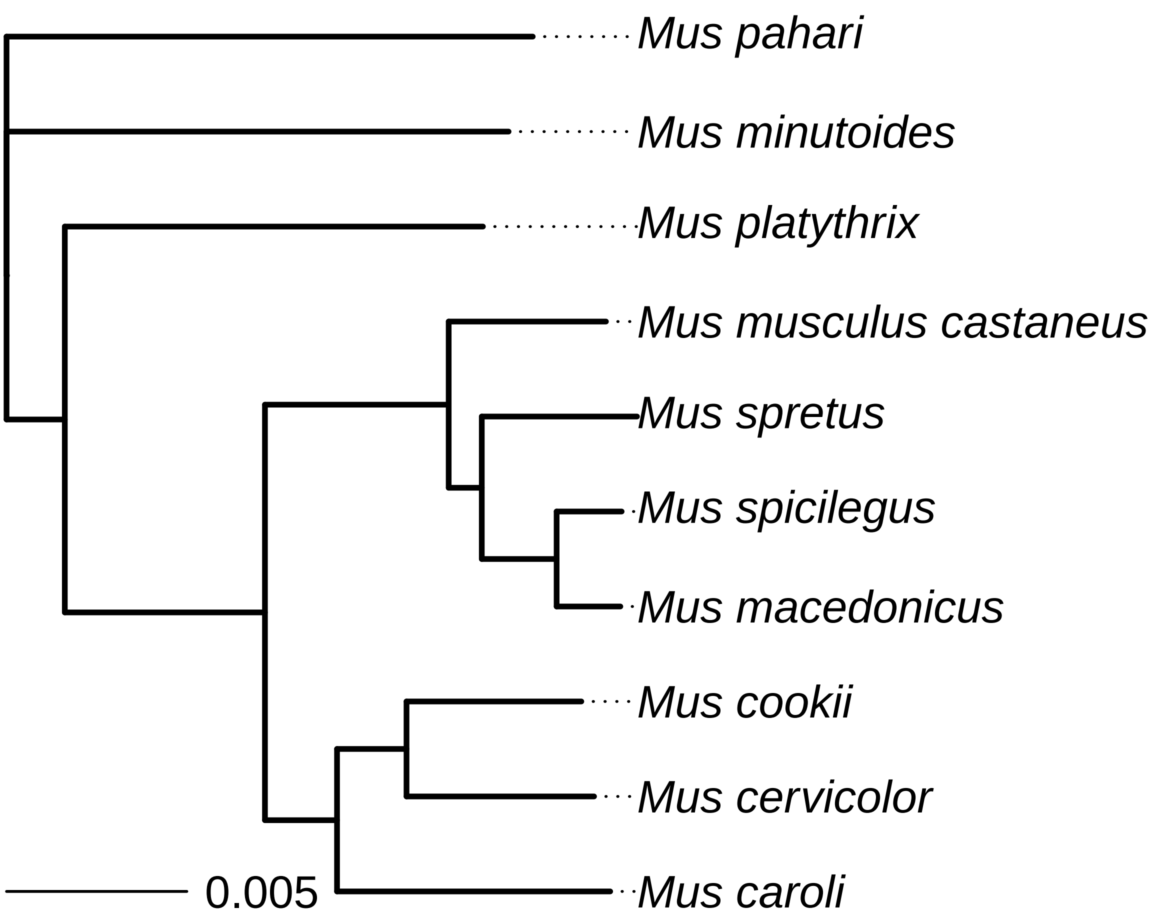


Figure S1: Phylogeny of the 10 *Murini* species, reconstructed from the concatenation of all exons. The scale is in substitution per site.

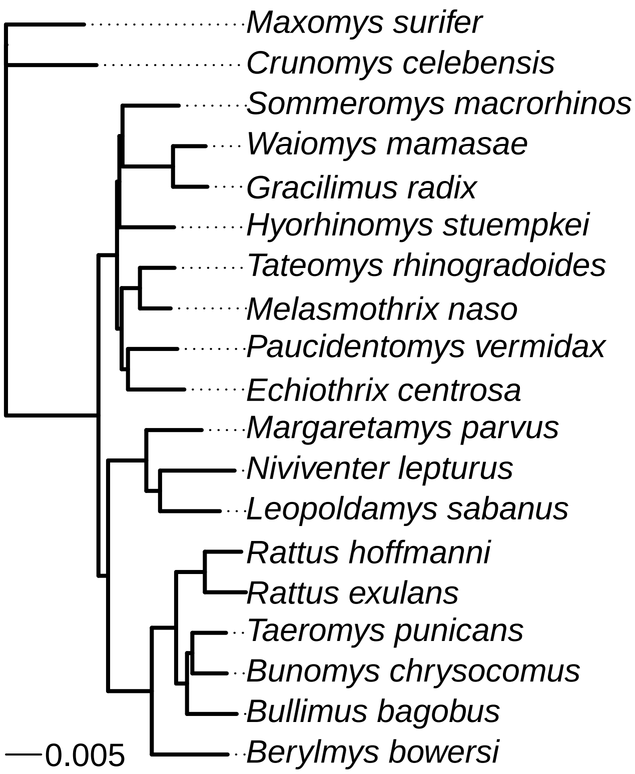


Figure S2: Phylogeny of the 19 *Rattini* species, reconstructed from the concatenation of all exons. The scale is in substitution per site.

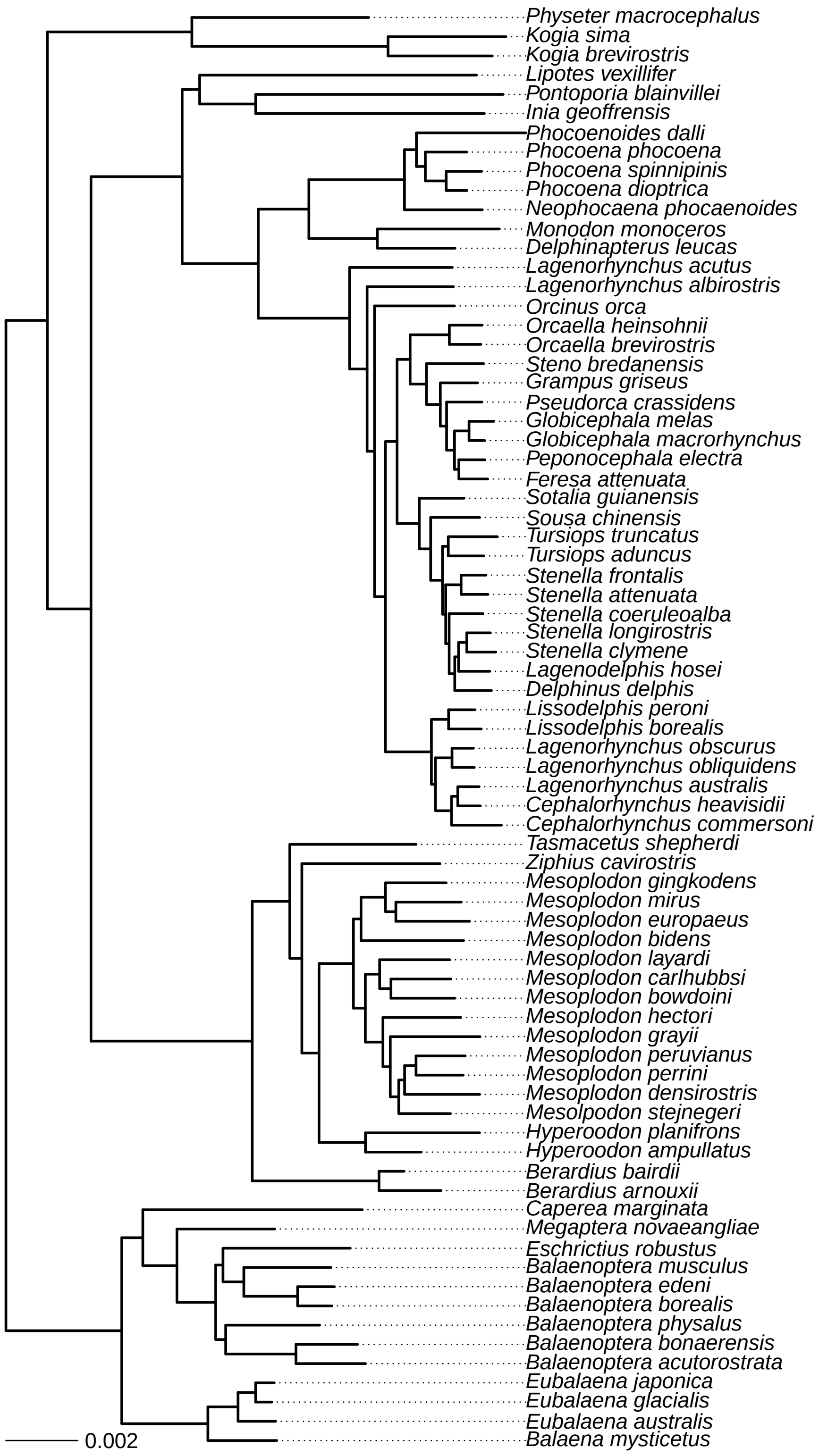


Figure S3: Phylogeny of the 75 *Cetacea* species, reconstructed from the concatenation of all exons. The scale is in substitution per site.

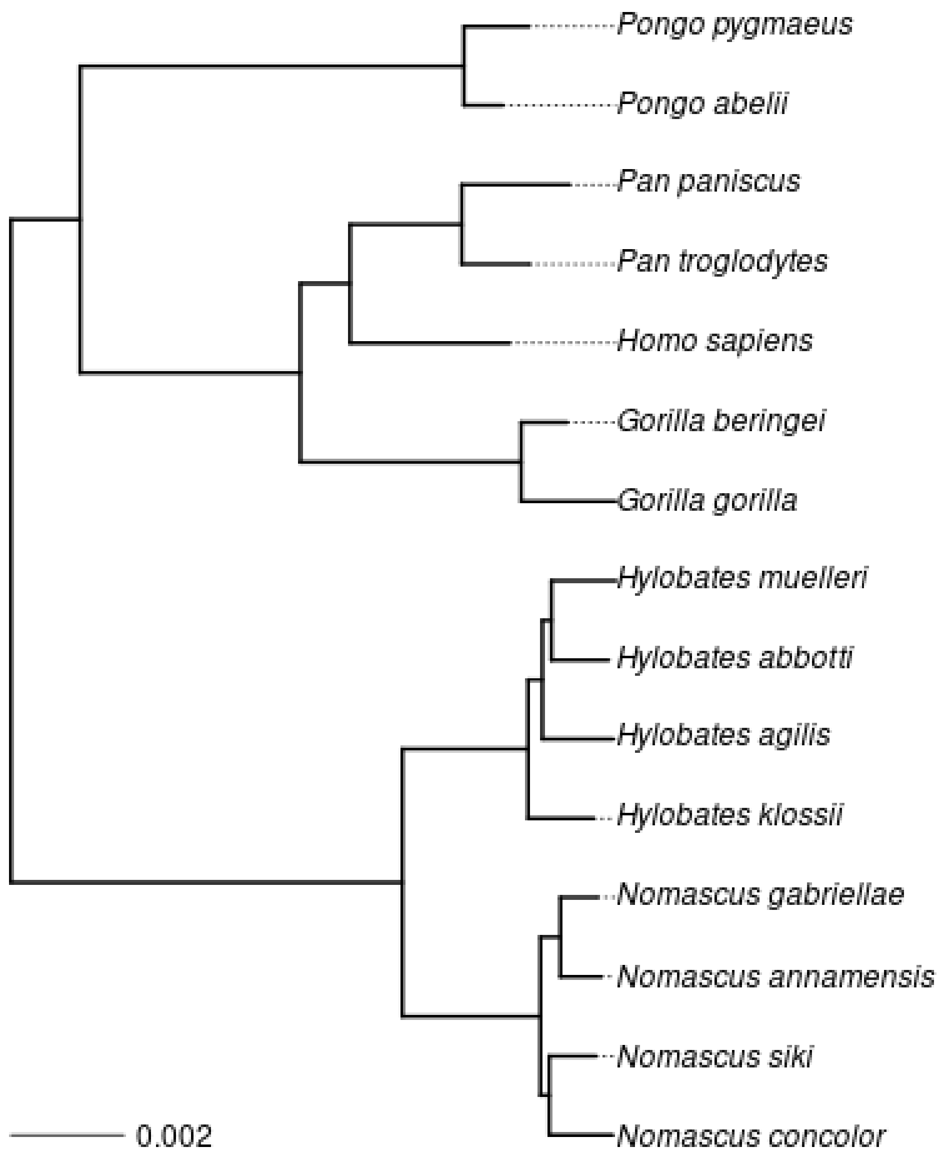


Figure S4: Phylogeny of the 15 *Hominoidae* species, reconstructed from the concatenation of all exons. The scale is in substitution per site.

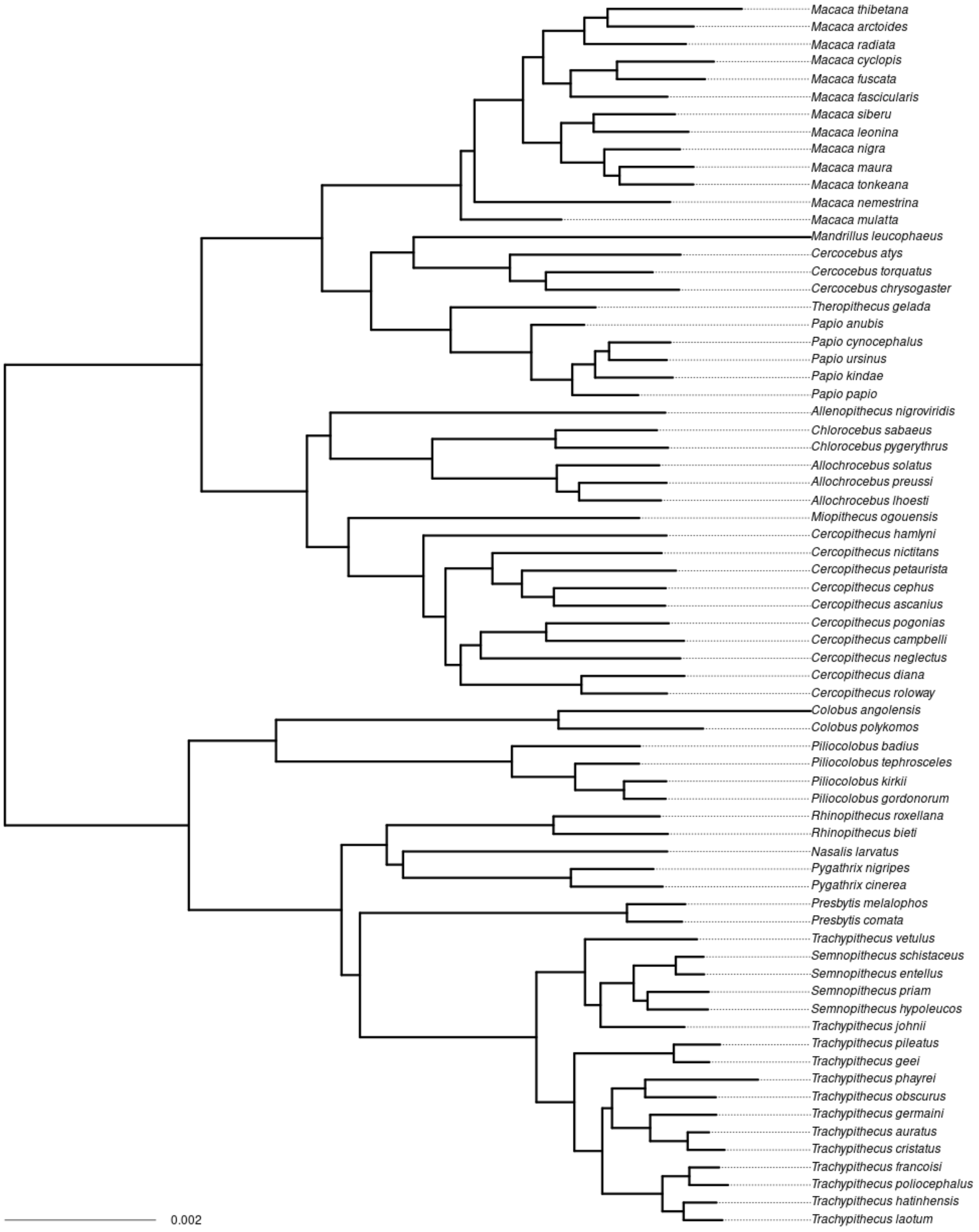


Figure S5: Phylogeny of the 70 *Cercopithecidae* species, reconstructed from the concatenation of all exons. The scale is in substitution per site.

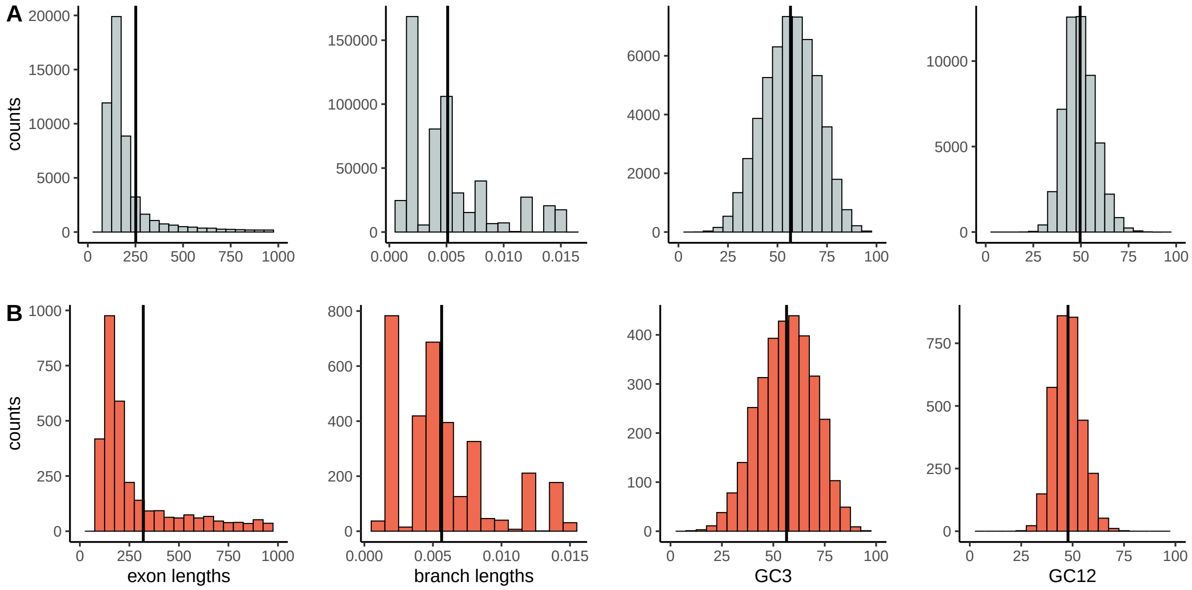


Figure S6: Distribution of exon length, branch length, GC3, GC12 in *Murini*. A: in all data, B: in gBGC episodes. Vertical lines represent the means.

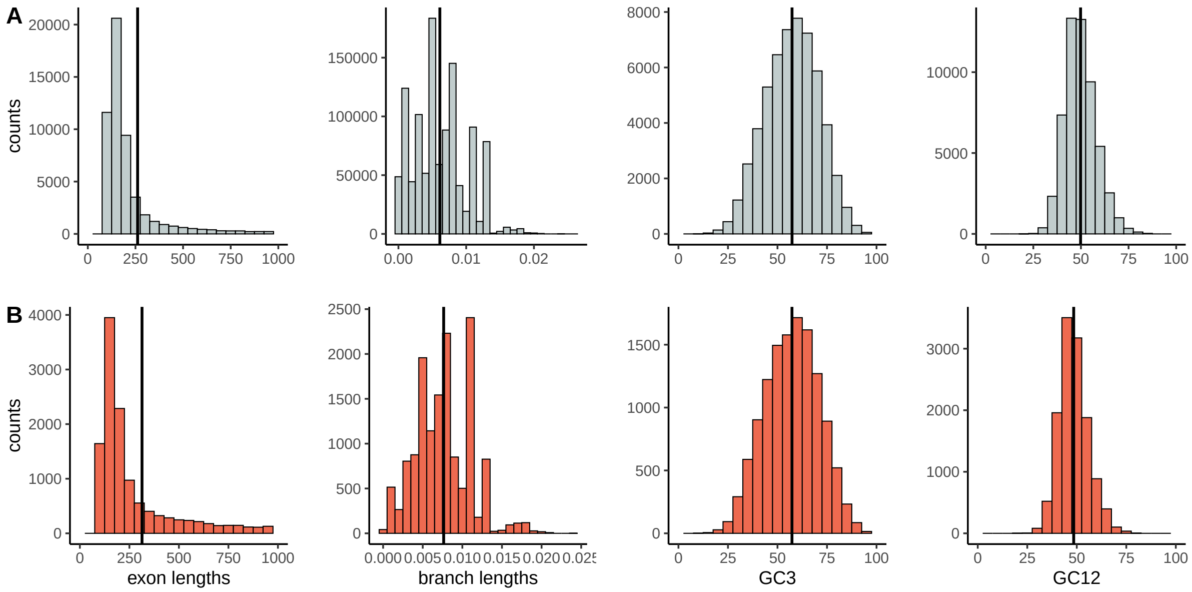


Figure S7: Distribution of exon length, branch length, GC3, GC12 in *Rattini*. A: in all data, B: in gBGC episodes. Vertical lines represent the means.

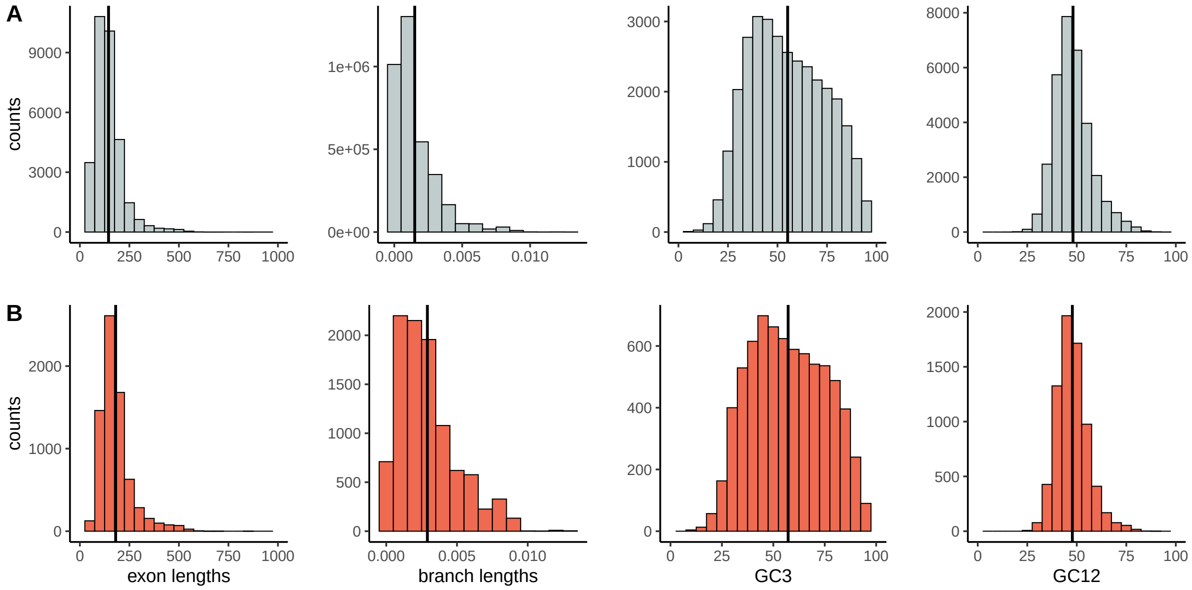


Figure S8: Distribution of exon length, branch length, GC3, GC12 in *Cetacea*. A: in all data, B: in gBGC episodes. Vertical lines represent the means.

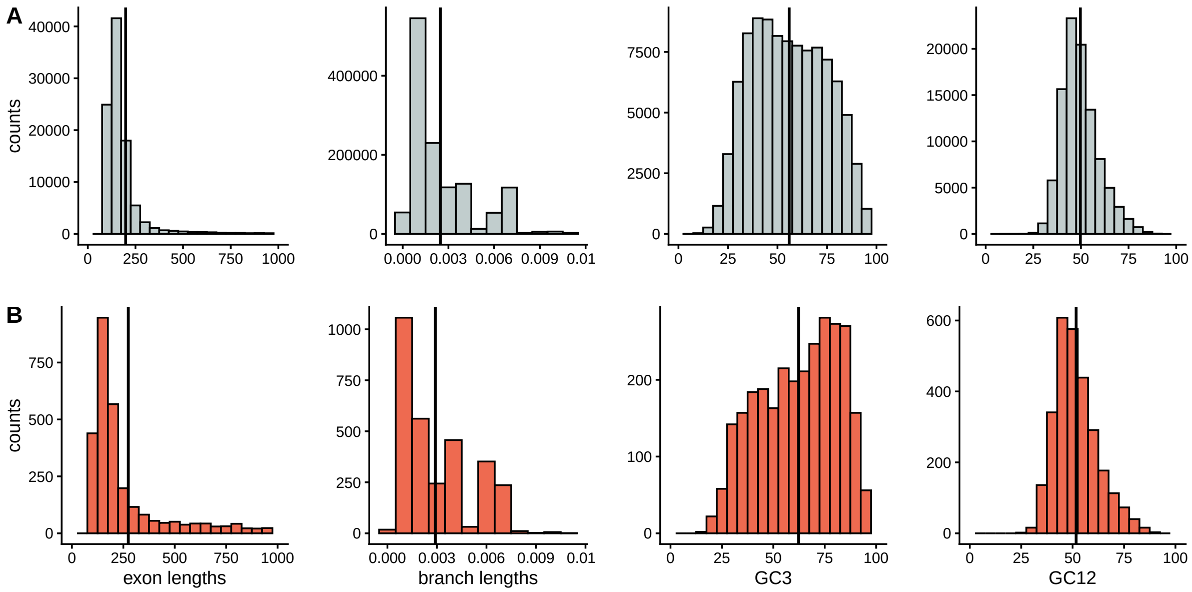


Figure S9: Distribution of exon length, branch length, GC3, GC12 in *Hominoidae*. A: in all data, B: in gBGC episodes. Vertical lines represent the means.

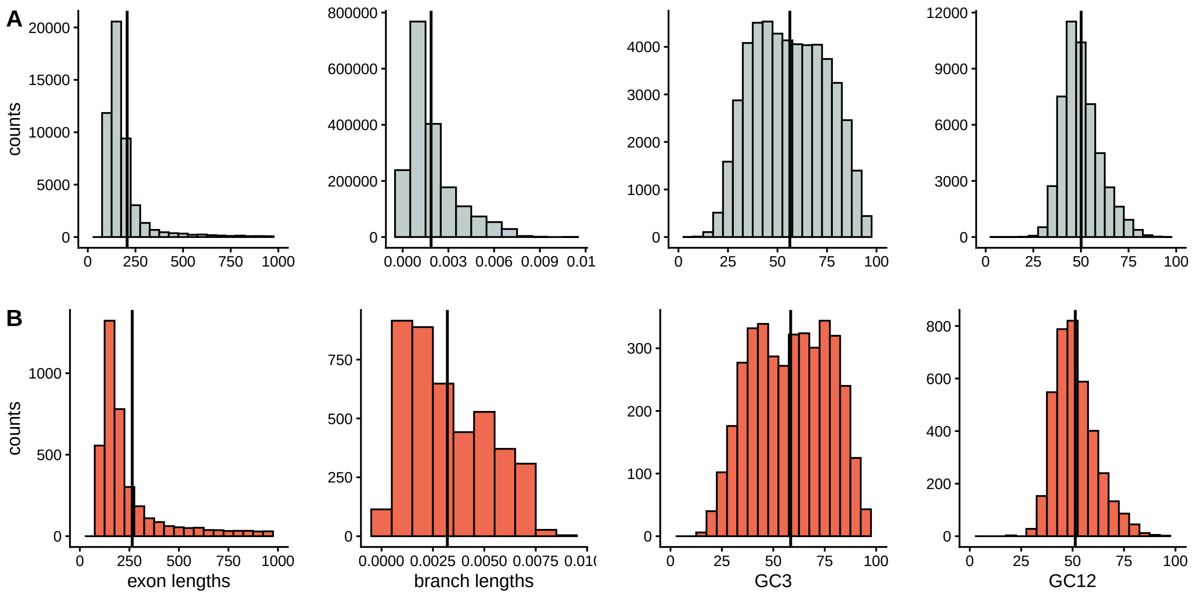


Figure S10: Distribution of exon length, branch length, GC3, GC12 in *Cercopithecidae*. A: in all data, B: in gBGC episodes. Vertical lines represent the means.

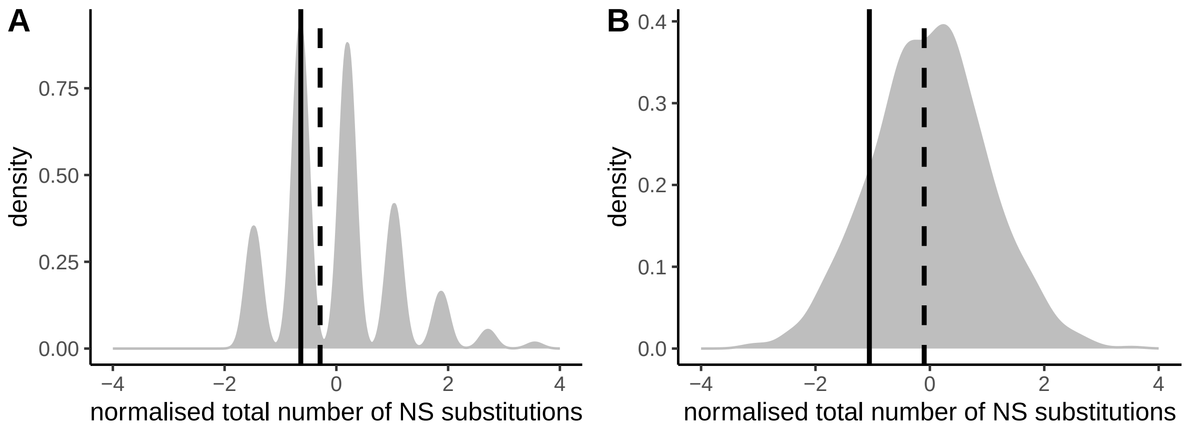


Figure S11: Protein evolution after punctual gBGC episodes in branches of length $\geq 0.006$ substitutions per site. A: *Murini*, B: *Rattini*. The vertical solid line corresponds to the observed number after deleterious episodes and the vertical dashed line corresponds to the observed number after non-deleterious episodes.

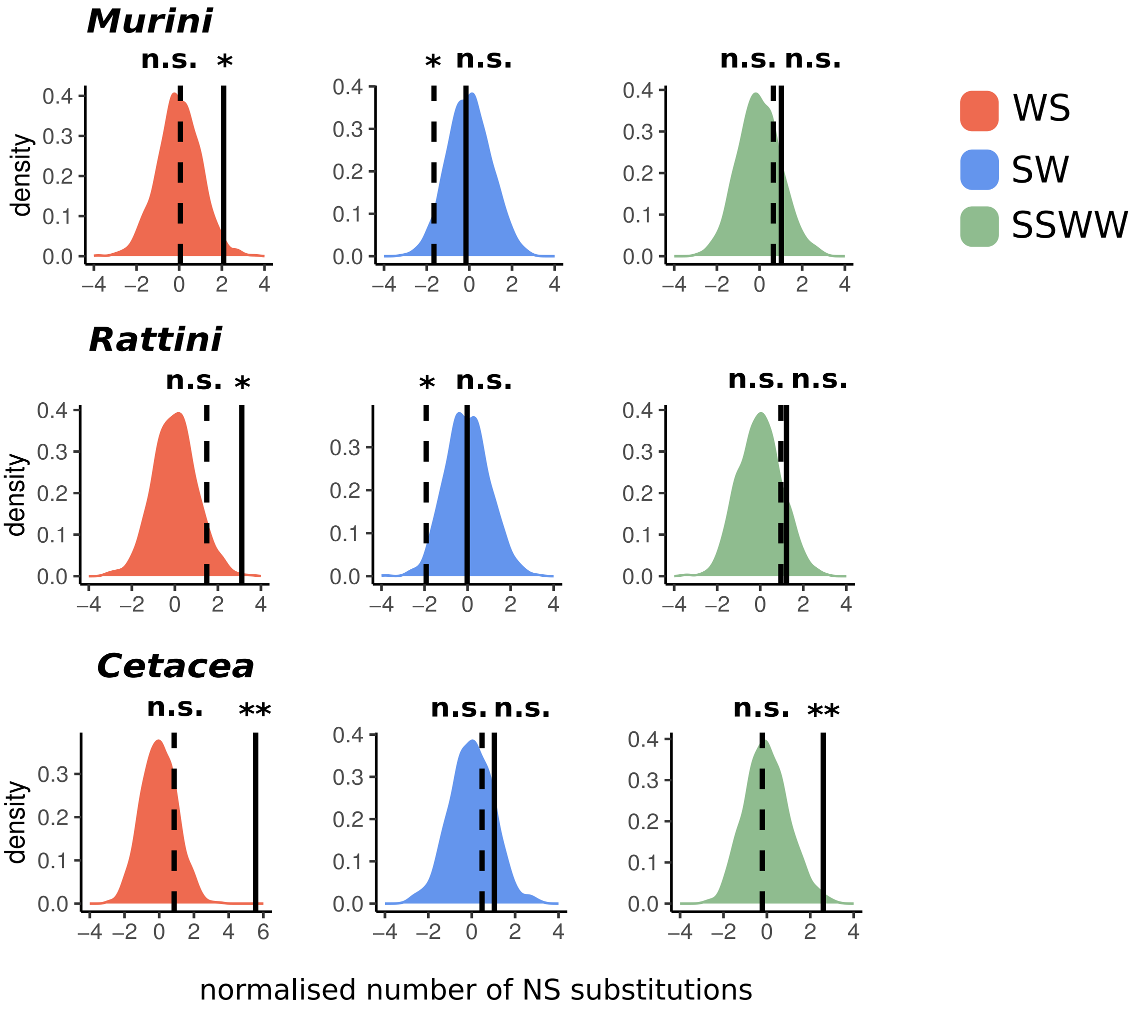


Figure S12: Response to punctual gBGC episodes detailed for each type of substitution, from left to right: WS, SW and SSWW. Results are shown for the adjusted branch lengths. The distribution shows the expected numbers from 1000 randomisations. The vertical solid line corresponds to the observed number after deleterious episodes and the vertical dashed line corresponds to the observed number after non-deleterious episodes. n.s. : p-value $\geq 0.05$, * : p-value $<$ 0.05, ** : p-value $<$ 0.01.

Table S1: Summary of the substitution mapping.

| Taxon | Substitution type | Total number in all data | Mean number in branches with punctual episodes | Mean number in branches without episodes | Ratio of mean number between branches with and without episodes | Mean number in branches after episodes (all types) | Mean number in branches after punctual episodes |
|---|---|---|---|---|---|---|---|
| **Murini** | S-WS | 155,214 | 3.00 | 0.22 | 13.7 | 0.33 | 0.26 |
| | S-SW | 230,734 | 0.46 | 0.27 | 1.7 | 0.39 | 0.32 |
| | S-SSWW | 34,731 | 0.16 | 0.046 | 3.5 | 0.068 | 0.056 |
| | NS-WS | 85,726 | 0.31 | 0.13 | 2.4 | 0.13 | 0.094 |
| | NS-SW | 123,851 | 0.15 | 0.14 | 1.1 | 0.15 | 0.10 |
| | NS-SSWW | 42,455 | 0.074 | 0.055 | 1.3 | 0.063 | 0.044 |
| **Rattini** | S-WS | 425,379 | 3.47 | 0.23 | 15.1 | 0.77 | 0.50 |
| | S-SW | 585,695 | 0.51 | 0.29 | 1.8 | 0.66 | 0.51 |
| | S-SSWW | 88,251 | 0.19 | 0.047 | 4.0 | 0.13 | 0.089 |
| | NS-WS | 242,615 | 0.52 | 0.13 | 4.0 | 0.32 | 0.19 |
| | NS-SW | 330,541 | 0.19 | 0.16 | 1.2 | 0.3 | 0.18 |
| | NS-SSWW | 112,471 | 0.14 | 0.056 | 2.5 | 0.13 | 0.071 |
| **Cetacea** | S-WS | 175,882 | 2.51 | 0.033 | 76.1 | 0.21 | 0.094 |
| | S-SW | 217,495 | 0.18 | 0.041 | 4.4 | 0.12 | 0.089 |
| | S-SSWW | 32,935 | 0.10 | 0.0064 | 15.6 | 0.031 | 0.021 |
| | NS-WS | 122,983 | 0.42 | 0.024 | 17.5 | 0.11 | 0.073 |
| | NS-SW | 130,411 | 0.13 | 0.024 | 5.4 | 0.058 | 0.050 |
| | NS-SSWW | 50,226 | 0.10 | 0.0093 | 10.8 | 0.033 | 0.027 |
| **Hominoidae** | S-WS | 144,576 | 2.53 | 0.14 | 18 .1 | 0.29 | 0.052 |
| | S-SW | 218,944 | 0.47 | 0.21 | 2.2 | 0.37 | 0.13 |
| | S-SSWW | 58,801 | 0.11 | 0.0034 | 32.4 | 0.25 | 0.017 |
| | NS-WS | 101,705 | 0.23 | 0.086 | 2.7 | 0.19 | 0.039 |
| | NS-SW | 139,432 | 0.23 | 0.11 | 2.1 | 0.21 | 0.058 |
| | NS-SSWW | 67,562 | 0.11 | 0.042 | 2.6 | 0.17 | 0.021 |
| **Cercopithecidae** | S-WS | 126,655 | 2.37 | 0.049 | 48.4 | 0.16 | 0.11 |
| | S-SW | 216,955 | 0.25 | 0.087 | 2.9 | 0.21 | 0.17 |
| | S-SSWW | 61,741 | 0.18 | 0.014 | 12.9 | 0.088 | 0.078 |
| | NS-WS | 97,413 | 0.38 | 0.032 | 11.9 | 0.15 | 0.11 |
| | NS-SW | 141,297 | 0.33 | 0.049 | 6.7 | 0.17 | 0.12 |
| | NS-SSWW | 72,603 | 0.22 | 0.017 | 12.9 | 0.12 | 0.072 |

Table S2: Composition and accession numbers of the rodents and primates datasets.

| Taxon | Species | Origin | Reference | Accession number |
|---|---|---|---|---|
| *Murini* | *Mus caroli* | NCBI | PRJNA323493 | SRR3601860,SRR3601849 |
| *Murini* | *Mus cervicolor* | NCBI | PRJNA323494 | SRR3601866,SRR3601865 |
| *Murini* | *Mus cookii* | NCBI | PRJNA323495 | SRR3601837,SRR3601836 |
| *Murini* | *Mus macedonicus* | NCBI | PRJNA323496 | SRR3601844,SRR3601842 |
| *Murini* | *Mus minutoides* | NCBI | PRJNA323497 | SRR3601846,SRR3601845 |
| *Murini* | *Mus musculus castaneus* | NCBI | PRJNA323498 | SRR3601864,SRR3601863 |
| *Murini* | *Mus pahari* | NCBI | PRJNA323499 | SRR3601853,SRR3601852 |
| *Murini* | *Mus platythrix* | NCBI | PRJNA323500 | SRR3601855,SRR3601854 |
| *Murini* | *Mus spicilegus* | NCBI | PRJNA323501 | SRR3601859,SRR3601858 |
| *Murini* | *Mus spretus* | NCBI | PRJNA323502 | SRR3601862,SRR3601861 |
| *Rattini* | *Berylmys bowersi* | NCBI | PRJNA705792 | SRR13848314 |
| *Rattini* | *Bullimus bagobus* | NCBI | PRJNA705793 | SRR13848281 |
| *Rattini* | *Bunomys chrysocomus* | NCBI | PRJNA705794 | SRR13848312 |
| *Rattini* | *Crunomys celebensis* | NCBI | PRJNA705795 | SRR13848291 |
| *Rattini* | *Echiothrix centrosa* | NCBI | PRJNA705796 | SRR13848287 |
| *Rattini* | *Hyorhinomys stuempkei* | NCBI | PRJNA705797 | SRR13848278 |
| *Rattini* | *Leopoldamys sabanus* | NCBI | PRJNA705798 | SRR13848288 |
| *Rattini* | *Margaretamys parvus* | NCBI | PRJNA705799 | SRR13848294 |
| *Rattini* | *Maxomys surifer* | NCBI | PRJNA705800 | SRR13848279 |
| *Rattini* | *Melasmothrix naso* | NCBI | PRJNA705801 | SRR13848293 |
| *Rattini* | *Gracilimus radix* | NCBI | PRJNA705802 | SRR13848298 |
| *Rattini* | *Niviventer lepturus* | NCBI | PRJNA705803 | SRR13848313 |
| *Rattini* | *Paucidentomys vermidax* | NCBI | PRJNA705804 | SRR13848290 |
| *Rattini* | *Rattus exulans* | NCBI | PRJNA705805 | SRR13848285 |
| *Rattini* | *Rattus hoffmanni* | NCBI | PRJNA705806 | SRR13848284 |
| *Rattini* | *Sommeromys macrorhinos* | NCBI | PRJNA705807 | SRR13848296 |
| *Rattini* | *Taeromys punicans* | NCBI | PRJNA705808 | SRR13848308 |
| *Rattini* | *Tateomys rhinogradoides* | NCBI | PRJNA705809 | SRR13848295 |
| *Rattini* | *Waiomys mamasae* | NCBI | PRJNA705810 | SRR13848297 |
| *Hominoidae* | *Pongo abelii* | Ensembl | Susie PABv2 release 113 | |
| *Hominoidae* | *Pongo pygmaeus* | NCBI | PRJNA785018 | ASM2376777v1 |
| *Hominoidae* | *Gorilla gorilla* | Ensembl | gorGor4 release 113 | |
| *Hominoidae* | *Gorilla beringei* | ENA | PRJEB67744 | CAUYTB010000000 |
| *Hominoidae* | *Homo sapiens* | Ensembl | GRCh38 release 112 | |
| *Hominoidae* | *Pan troglodytes* | Ensembl | Pan tro 3 release 112 | |
| *Hominoidae* | *Pan paniscus* | Ensembl | Panpan1.1 release 113 | |

| Taxon | Species | Origin | Reference | Accession number |
|---|---|---|---|---|
| *Hominoidae* | *Nomascus concolor* | ENA | PRJEB67744 | CAUYSP010000000 |
| *Hominoidae* | *Nomascus siki* | ENA | PRJEB67744 | CAUYNF010000000 |
| *Hominoidae* | *Nomascus annamensis* | ENA | PRJEB67744 | CAUYQL010000000 |
| *Hominoidae* | *Nomascus gabriellae* | ENA | PRJEB67744 | CAUYQO010000000 |
| *Hominoidae* | *Hylobates klossii* | ENA | PRJEB67744 | CAUYPF010000000 |
| *Hominoidae* | *Hylobates agilis* | ENA | PRJEB67744 | CAUYOT010000000 |
| *Hominoidae* | *Hylobates abbotti* | ENA | PRJEB67744 | CAUYSE010000000 |
| *Hominoidae* | *Hylobates muelleri* | ENA | PRJEB67744 | CAUYNX010000000 |
| *Cercopithecidae* | *Macaca arctoides* | ENA | PRJEB67744 | CAUYNL010000000 |
| *Cercopithecidae* | *Macaca tonkeana* | ENA | PRJEB67744 | CAUYNJ010000000 |
| *Cercopithecidae* | *Macaca nigra* | ENA | PRJEB67744 | CAUYNM010000000 |
| *Cercopithecidae* | *Macaca radiata* | ENA | PRJEB67744 | CAUYNO010000000 |
| *Cercopithecidae* | *Macaca leonina* | ENA | PRJEB67744 | CAUYNU010000000 |
| *Cercopithecidae* | *Macaca siberu* | ENA | PRJEB67744 | CAUYPE010000000 |
| *Cercopithecidae* | *Macaca fascicularis* | ENA | PRJEB67744 | CAUYPJ010000000 |
| *Cercopithecidae* | *Macaca fuscata* | ENA | PRJEB67744 | CAUYPW010000000 |
| *Cercopithecidae* | *Macaca cyclopis* | ENA | PRJEB67744 | CAUYPY010000000 |
| *Cercopithecidae* | *Macaca thibetana* | ENA | PRJEB67744 | CAUYQE010000000 |
| *Cercopithecidae* | *Macaca maura* | ENA | PRJEB67744 | CAUYSB010000000 |
| *Cercopithecidae* | *Macaca mulatta* | Ensembl | Mmul_10 release 113 | |
| *Cercopithecidae* | *Papio papio* | ENA | PRJEB67744 | CAUYMZ010000000 |
| *Cercopithecidae* | *Papio kindae* | ENA | PRJEB67744 | CAUYNZ010000000 |
| *Cercopithecidae* | *Papio ursinus* | ENA | PRJEB67744 | CAUYQN010000000 |
| *Cercopithecidae* | *Papio cynocephalus* | ENA | PRJEB67744 | CAUYRL010000000 |
| *Cercopithecidae* | *Cercocebus chrysogaster* | ENA | PRJEB67744 | CAUYNG010000000 |
| *Cercopithecidae* | *Cercocebus torquatus* | ENA | PRJEB67744 | CAUYOL010000000 |
| *Cercopithecidae* | *Cercopithecus roloway* | ENA | PRJEB67744 | CAUYNT010000000 |
| *Cercopithecidae* | *Cercopithecus ascanius* | ENA | PRJEB67744 | CAUYPR010000000 |
| *Cercopithecidae* | *Cercopithecus neglectus* | ENA | PRJEB67744 | CAUYQZ010000000 |
| *Cercopithecidae* | *Cercopithecus campbelli* | ENA | PRJEB67744 | CAUYRC010000000 |
| *Cercopithecidae* | *Cercopithecus hamlyni* | ENA | PRJEB67744 | CAUYRU010000000 |
| *Cercopithecidae* | *Cercopithecus diana* | ENA | PRJEB67744 | CAUYRX010000000 |
| *Cercopithecidae* | *Cercopithecus pogonias* | ENA | PRJEB67744 | CAUYSS010000000 |
| *Cercopithecidae* | *Cercopithecus nictitans* | ENA | PRJEB67744 | CAUYSU010000000 |
| *Cercopithecidae* | *Cercopithecus petaurista* | ENA | PRJEB67744 | CAUYSW010000000 |
| *Cercopithecidae* | *Cercopithecus cephus* | ENA | PRJEB67744 | CAUYUF010000000 |
| *Cercopithecidae* | *Miopithecus ogouensis* | ENA | PRJEB67744 | CAUYND010000000 |
| *Cercopithecidae* | *Allochrocebus lhoesti* | ENA | PRJEB67744 | CAUYOZ010000000 |

| Taxon | Species | Origin | Reference | Accession number |
|---|---|---|---|---|
| *Cercopithecidae* | *Allochrocebus preussi* | ENA | PRJEB67744 | CAUYSH010000000 |
| *Cercopithecidae* | *Allochrocebus solatus* | ENA | PRJEB67744 | CAUYTQ010000000 |
| *Cercopithecidae* | *Chlorocebus pygerythrus* | ENA | PRJEB67744 | CAUYTG010000000 |
| *Cercopithecidae* | *Allenopithecus nigroviridis* | ENA | PRJEB67744 | CAUYOP010000000 |
| *Cercopithecidae* | *Trachypithecus laotum* | ENA | PRJEB67744 | CAUYNP010000000 |
| *Cercopithecidae* | *Trachypithecus geei* | ENA | PRJEB67744 | CAUYNV010000000 |
| *Cercopithecidae* | *Trachypithecus hatinhensis* | ENA | PRJEB67744 | CAUYOF010000000 |
| *Cercopithecidae* | *Trachypithecus cristatus* | ENA | PRJEB67744 | CAUYOG010000000 |
| *Cercopithecidae* | *Trachypithecus germaini* | ENA | PRJEB67744 | CAUYOH010000000 |
| *Cercopithecidae* | *Trachypithecus pileatus* | ENA | PRJEB67744 | CAUYOM010000000 |
| *Cercopithecidae* | *Trachypithecus obscurus* | ENA | PRJEB67744 | CAUYOO010000000 |
| *Cercopithecidae* | *Trachypithecus phayrei* | ENA | PRJEB67744 | CAUYOV010000000 |
| *Cercopithecidae* | *Trachypithecus auratus* | ENA | PRJEB67744 | CAUYPD010000000 |
| *Cercopithecidae* | *Trachypithecus johnii* | ENA | PRJEB67744 | CAUYPG010000000 |
| *Cercopithecidae* | *Trachypithecus poliocephalus* | ENA | PRJEB67744 | CAUYPM010000000 |
| *Cercopithecidae* | *Trachypithecus phayrei* | ENA | PRJEB67744 | CAUYQF010000000 |
| *Cercopithecidae* | *Trachypithecus francoisi* | ENA | PRJEB67744 | CAUYRV010000000 |
| *Cercopithecidae* | *Trachypithecus vetulus* | ENA | PRJEB67744 | CAUYST010000000 |
| *Cercopithecidae* | *Semnopithecus hypoleucos* | ENA | PRJEB67744 | CAUYNQ010000000 |
| *Cercopithecidae* | *Semnopithecus entellus* | ENA | PRJEB67744 | CAUYOJ010000000 |
| *Cercopithecidae* | *Semnopithecus priam* | ENA | PRJEB67744 | CAUYPN010000000 |
| *Cercopithecidae* | *Semnopithecus schistaceus* | ENA | PRJEB67744 | CAUYPV010000000 |
| *Cercopithecidae* | *Pygathrix cinerea* | ENA | PRJEB67744 | CAUYNW010000000 |
| *Cercopithecidae* | *Pygathrix nigripes* | ENA | PRJEB67744 | CAUYOA010000000 |
| *Cercopithecidae* | *Rhinopithecus bieti* | ENA | PRJEB67744 | CAUYQC010000000 |
| *Cercopithecidae* | *Nasalis larvatus* | ENA | PRJEB67744 | CAUYSA010000000 |
| *Cercopithecidae* | *Presbytis comata* | ENA | PRJEB67744 | CAUYOK010000000 |
| *Cercopithecidae* | *Presbytis melalophos* | ENA | PRJEB67744 | CAUYTV010000000 |
| *Cercopithecidae* | *Piliocolobus gordonorum* | ENA | PRJEB67744 | CAUYPL010000000 |
| *Cercopithecidae* | *Piliocolobus badius* | ENA | PRJEB67744 | CAUYPZ010000000 |
| *Cercopithecidae* | *Piliocolobus kirkii* | ENA | PRJEB67744 | CAUYRS010000000 |
| *Cercopithecidae* | *Colobus polykomos* | ENA | PRJEB67744 | CAUYRQ010000000 |
| *Cercopithecidae* | *Colobus angolensis* | Ensembl | Cang.pa_1.0 release 114 | |
| *Cercopithecidae* | *Mandrillus leucophaeus* | Ensembl | Mleu.le_1.0 release 114 | |
| *Cercopithecidae* | *Theropithecus gelada* | Ensembl | Tgel_1.0 release 114 | |
| *Cercopithecidae* | *Rhinopithecus roxellana* | Ensembl | Rrox_v1 release 114 | |
| *Cercopithecidae* | *Papio anubis* | Ensembl | Panubis1.0 release 114 | |
| *Cercopithecidae* | *Macaca nemestrina* | Ensembl | Mnem_1.0 release 114 | |

| Taxon | Species | Origin | Reference | Accession number |
|---|---|---|---|---|
| *Cercopithecidae* | *Cercocebus atys* | Ensembl | Caty_1.0 release 114 | |
| *Cercopithecidae* | *Piliocolobus tephrosceles* | Ensembl | ASM277652v2 release 114 | |
| *Cercopithecidae* | *Chlorocebus sabaeus* | Ensembl | ChlSab1.1 release 114 | |
